# DNA polymerase I is an efficient reverse transcriptase that mediates RNA-templated DNA repair synthesis

**DOI:** 10.64898/2025.12.22.696026

**Authors:** Frances C. Lowder, Abigail H. Kendal, Lyle A. Simmons

## Abstract

It is well established that during normal growth and under stress conditions ribonucleotides become nested in genomic DNA. Given the high frequency with which RNA is found in genomic DNA, it is unknown which DNA polymerase is responsible for catalyzing RNA-dependent DNA synthesis. In this work we establish bacterial DNA polymerase I (Pol I) as a robust reverse transcriptase. We tested several bacterial Pol I enzymes and found that all possess RT activity with *B. subtilis* Pol I showing RNA-dependent DNA synthesis activity similar to the Moloney Murine Leukemia Virus reverse transcriptase. We tested RNA embedded in DNA and found that replicative DNA polymerase PolC arrests at a single template ribonucleotide while, in contrast, Pol I is able to efficiently traverse long stretches of templated ribonucleotides. We conclude that Pol I is responsible for RNA templated DNA replication, allowing for replication forks to navigate RNA that persists in genomic DNA.

## Introduction

RNA-DNA hybrids represent one of the most prevalent and pervasive endogenous nucleic acids that leads to genome instability in all organisms (summarized in Figure 1A, reviewed in^1^). Although RNA-DNA hybrids are well documented for their detrimental effects on genome integrity, hybrid formation is essential for all cells due to the role of hybrids in priming DNA synthesis^2,3^ and in the production of mRNA during transcription^4,5^. RNA-DNA hybrids also contribute to DNA repair during sub-optimal conditions^6,7^ and regulate gene expression^8^. Failure to resolve hybrids is a driving force for genome instability in organisms ranging from bacteria to humans and underlies the development of several human cancers and neurological disorders^9,10^. The nucleolytic activity of the 2′-OH of the ribose sugars can nick the phosphodiester backbone of nearby DNA, leading to the formation of DNA strand breaks^11^. Persistent RNA-DNA hybrids resulting from RNA polymerase impose barriers to DNA replication causing fork stalling^12–15^, induction of the DNA damage response^16^, hyper-recombination and genome rearrangements^17,18^, all of which threaten cell survival, fitness or have the potential to lead to the development of human diseases^19,20^.

**Figure 1.**
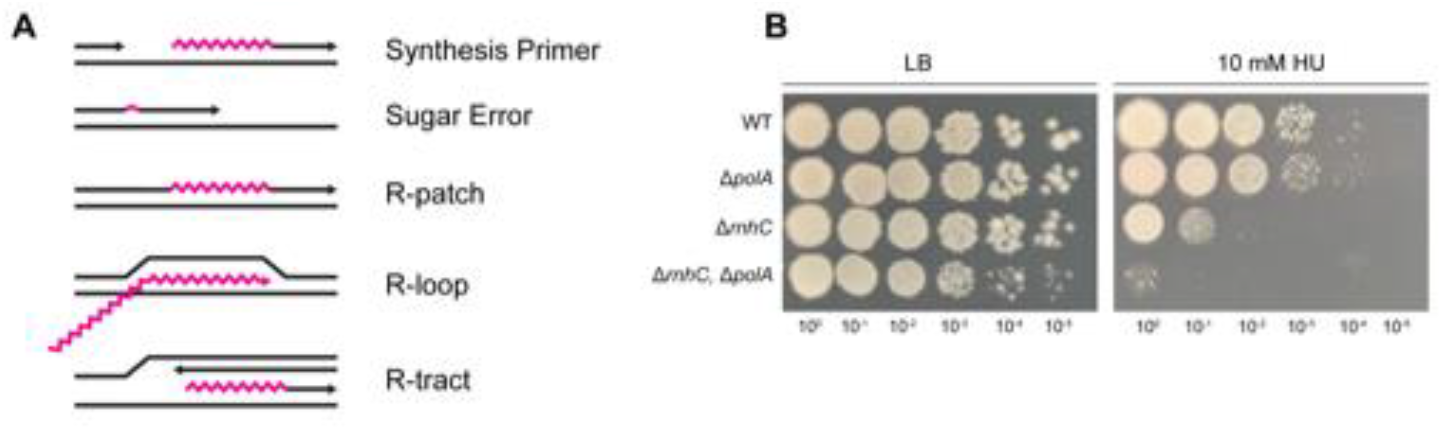
The absence of BsPol I sensitizes cells to ribonucleotide incorporation. (A) Depictions of the major types of RNA-DNA hybrids. (B) Spot titers of deletion strains treated with hydroxyurea (HU).

It is well established that replicative DNA polymerases from bacteria to eukaryotes misincorporate ribonucleotides (rNMPs) in place of their cognate deoxyribonucleotides^21–25^. Sugar errors are a major source of RNA-DNA hybrids that become embedded in the genome, as rNMP errors occur on the order of a couple thousand per round of replication for a bacterium^24^ to over a million per round of replication for mammals^21,26^. Should these ribonucleotide errors persist in the template strand, they slow or arrest progression of replicative bacterial DNA polymerase III^24^ and eukaryotic DNA polymerase δ^27,28^. In addition to single ribonucleotide errors, stretches of 4 or more rNMPs can be incorporated into genomic DNA. These include R-patches, which arise from errors by translesion DNA polymerases^29^, and R-tracts, which result from the integration of R-loops formed during transcription^30^. These extended stretches of RNA embedded in DNA pose significant problems for DNA replication^24,26,27^. When left unrepaired, cells would need to use a reverse transcriptase to catalyze RNA-templated DNA synthesis across the embedded RNA or the replication fork would arrest^31^. Failure to bypass or repair the embedded ribonucleotides would cause fork demise and activation of the DNA damage response, the mutagenic consequences of which threaten genome integrity^32^.

A-family DNA polymerases are well represented among bacteria, viruses, bacteriophages and eukaryotes. The archetypal A-family DNA polymerase is *Escherichia coli* DNA polymerase I (Pol I)^33^. In addition to DNA synthesis activity, *E. coli* Pol I has 5′ to 3′ exonuclease activity and 3′ to 5′ proofreading exonuclease activity, and as such it performs high fidelity DNA synthesis^34–36^.

Furthermore, it has been shown that a modified version of the *E. coli* Pol I Klenow lacking the 3′ to 5′ proofreading exonuclease activity has reverse transcriptase activity^37^. The eukaryotic DNA polymerase, Pol q, is able to traverse rNMPs embedded in template DNA^38,39^. Pol q is another A-family DNA polymerase, that has an inactive proofreading domain^38,39^, is highly error prone and has also been shown to catalyze RNA-templated DNA repair^37^. Work by Chandramouly et al. suggests that A-family DNA polymerases lacking proofreading activity may also have innate reverse transcriptase activity that would be important for RNA-templated DNA repair^37^. This link between error prone A-family DNA polymerases and reverse transcriptase activity is further bolstered by observations that Pol I enzymes from thermophilic bacteria that lack 3′ to 5′ proofreading activity have also been shown to possess some reverse transcriptase activity^40^.

As there are numerous studies of *E. coli* Pol I, several of which are foundational to our understanding of DNA replication, this protein has long been the model for bacterial Pol I enzymes^33^. *E. coli*, however, is a Gram-negative bacterium; in contrast, many Gram-positive bacteria have evolved a Pol I enzyme that lacks 3′ to 5′ proofreading exonuclease activity, due to mutations that prevent coordination of the Mg^2+^ required for catalysis^41,42^. This vestigial proofreading domain of Gram-positive Pol I is reminiscent of the inactive proofreading domain of eukaryotic Pol q which allows for Pol q to become error prone and may allow for RNA-templated DNA repair^39,43^. Consequently, we hypothesized that Gram-positive Pol I enzymes may have innate reverse transcriptase activity and could therefore catalyze RNA-templated DNA repair over unrepaired ribonucleotides that are embedded in the genome.

Here, we investigated the impact of ribonucleotide-containing templates on the DNA polymerase activity of A and C-family DNA polymerases from the Gram-positive bacterium *Bacillus subtilis*. We found that the presence of RNA in the template strand rapidly arrests synthesis by replicative C-family DNA polymerases PolC and DnaE. In contrast, we found that *B. subtilis* Pol I (*Bs*Pol I) is capable of robust reverse transcriptase activity, demonstrating the potential for RNA-templated DNA repair activity by *Bs*Pol I. We show that Pol I traverses DNA templates containing consecutive ribonucleotides, demonstrating that Pol I is able to replace a replicative DNA polymerase when a ribonucleotide patch is encountered. We expanded our findings and tested several other bacterial Pol I enzymes and found that many have reverse transcriptase activity, albeit with varying efficiencies. Finally, we demonstrate that ribonucleotide incorporation in DNA represents a source of genome instability that is exacerbated in the absence of Pol I. Together, this study demonstrates the critical contribution of RNA-templated DNA repair activity by bacterial Pol I, while also establishing Pol I as a robust reverse transcriptase that is important for genome stability by allowing replication across ribonucleotides embedded in genomic DNA.

## Results

### The absence of *Bs*Pol I sensitizes cells to ribonucleotide incorporation

Although there is substantial evidence that ribonucleotides are embedded in genomic DNA^28,29,44–48^, we asked whether a loss of Pol I sensitized cells to increased ribonucleotide concentrations in vivo. To explore this question, we challenged cells with hydroxyurea (HU), a compound that inhibits ribonucleotide reductase and increases the occurrence of RNA-DNA hybrids, particularly sugar errors and R-patches (Figure 1A)^49^. As shown in Figure 1B, Δ*polA* strains are mildly sensitive to HU, showing reduced growth at 10 mM HU. Cells lacking Ribonuclease HIII (Δ*rnhC*, RNase HIII) display a more severe growth phenotype on HU, owing to the role of this enzyme in resolving RNA-DNA hybrids^50–52^. Importantly, loss of *rnhC* is known to increase the occurrence of R-loops in *B. subtilis*^52^. The cells that lack both Pol I and RNase HIII are most sensitive to replication stress caused by HU. This result emphasizes the importance of Pol I responding to RNA-DNA hybrids in vivo. Therefore, the sensitivity of the Δ*polA* strain is in part caused by a failure of replication over RNA nested in the template stand.

### *Bacillus subtilis* Pol I can use RNA as a synthesis template

While Pol I is strongly associated with DNA repair, we were also interested in testing several DNA polymerases for RNA templated DNA synthesis. Because *polC* and *dnaE* are essential genes^53,54^, we used a biochemical approach to determine if replicative DNA polymerases could synthesize DNA using a template comprised entirely of ribonucleotides. Using purified Pol I, PolC, and DnaE (Sup. Fig. 1), we assayed primer extension from a DNA or an RNA template. As a control, we also assayed the known reverse transcriptase (RT) from Moloney Murine Leukemia Virus (M-MuLV)^55^. Figure 2A demonstrates that all three *B. subtilis* polymerases were catalytically active, with robust activity using a DNA template in a buffer optimized for *B. subtilis* replication^53^ (Ext. Buffer). All three polymerases were also active in a simpler buffer used to promote RT activity (RT Buffer), however there were minor differences in the final products generated (Fig. 2B). Note that the RNA template migration is slower than the DNA product as slower RNA migration is attributed to the properties of RNA relative to DNA. When the template strand was composed entirely of RNA, Pol I was the only DNA polymerase able to fully traverse RNA and generate the same product as the RT control in either buffer (magenta band) (Fig. 2C and D). Conversely, PolC produced consistent intermediate products using the RNA template, while synthesis by DnaE was completely blocked (Fig. 2C and D). If the RNA template was twice as long, Pol I still produced similar products to the control RT while PolC and DnaE could only perform minimal primer extension (Fig. 2E and F). We conclude that wild type Pol I from *B. subtilis* is a robust reverse transcriptase, and that replicative C-family polymerases are unable to engage in significant RNA-templated DNA synthesis. We note that Pol I can perform RNA-templated DNA synthesis under different assay conditions, suggesting that RT activity is not dependent on a specific in vitro experimental design representing an authentic activity of Pol I.

**Figure 2.**
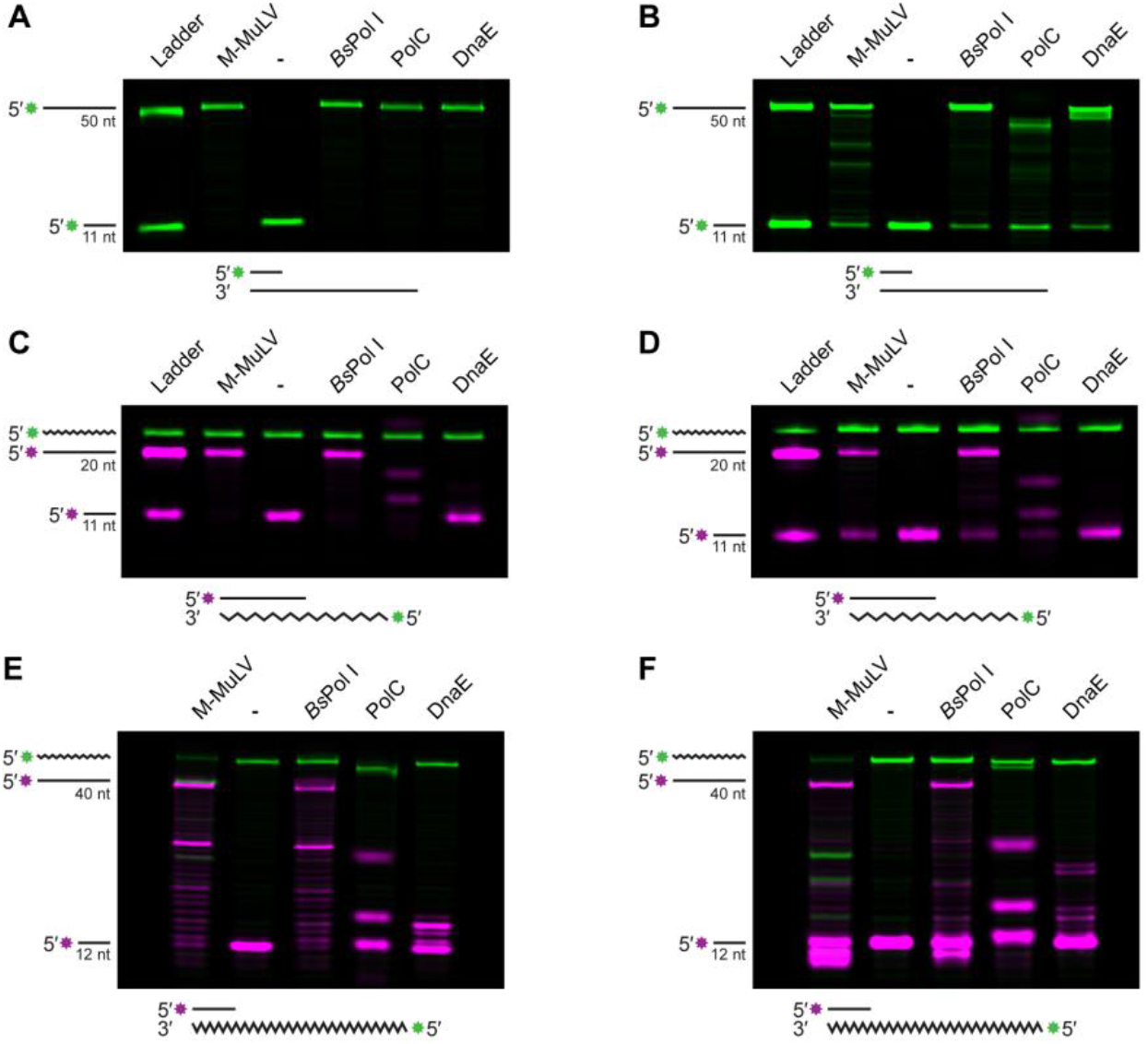
BsPol I can use RNA as a synthesis template. (A-C) Primer extension using a 50 nt DNA template (A), a 20 nt RNA template (B) or a 40 nt RNA template using a buffer optimized for extension. (D-F) The same as A-C, except reactions used a buffer optimized for reverse-transcriptase activity.

### RNA-templated DNA repair is conserved in bacterial Pol I enzymes

Based on our biochemical results, we asked if the RT activity we observed for *Bs*PoI I was conserved among other bacterial Pol I enzymes. We compiled a list of other instances where Pol I enzymes had been shown to have RT activity (Sup. Table 1). Given the variety of experimental conditions in these works, no clear comparison can be made from this data, however several trends are noted. Of the 22 bacterial Pol Is shown to have RT activity, only 5 have all four residues of the DEDD motif, which is responsible for coordination of the metals needed for 3′ to 5′ exonuclease activity^56^. These species also maintain the same base-stacking residue, a tyrosine that helps orient the nucleotide at the 3′ end of the substrate^56^. The remaining species are predicted to have catalytically inactive 3′-5′ exonuclease domains and have a mix of tyrosine or histidine residues at the base-stacking position (Sup. Table 1). Strikingly, the majority of Pol Is studied originate from extremophiles. As these proteins are stable at higher temperatures, the investigation of these Pol Is are most relevant for commercial applications necessitating specific in vitro conditions. To understand the conservation of Pol I RT activity, we chose to explore versions of this enzyme from model organisms and bacterial pathogens, with a focus on proteins with catalytically inactive proofreading domains since Pol q^37^ and now *B. subtilis* Pol I are adept reverse transcriptases.

We assayed a selection of Pol Is from *E. coli, S. aureus, M. smegmatis* and *L. monocytogenes*. All four proteins were expressed and purified from *E. coli* (Sup. Fig. 1) and assayed using the same DNA-DNA and RNA-DNA templates as Figure 2. The DNA polymerase activity of these proteins was confirmed using a DNA template, where all proteins show successful primer extension in either assay condition (Fig. 3A and B). When given an RNA template, the RT activities of the proteins varied. In the DNA polymerase Buffer, *Bs*Pol I and *Sa*Pol I produced full-length product, while and *Ec*Pol I produced some full-length product (Fig. 3C). *Lm*Pol I and *Ms*Pol I produced small amounts of fully synthesized DNA in this buffer, with their major products stalling about halfway through synthesis (Fig. 3C). When the assays were performed using the RT-optimized buffer, all four polymerases were able to generate full-length products (Fig. 3D). When a longer RNA template was used, the results were similar (Fig. 3E and F). *Sa*Pol I and *Ec*Pol I behaved more similarly to *Bs*Pol I than *Lm*Pol and *Ms*Pol I, but *Bs*Pol I consistently had fewer intermediate products remaining after 60 minutes of extension.

**Figure 3.**
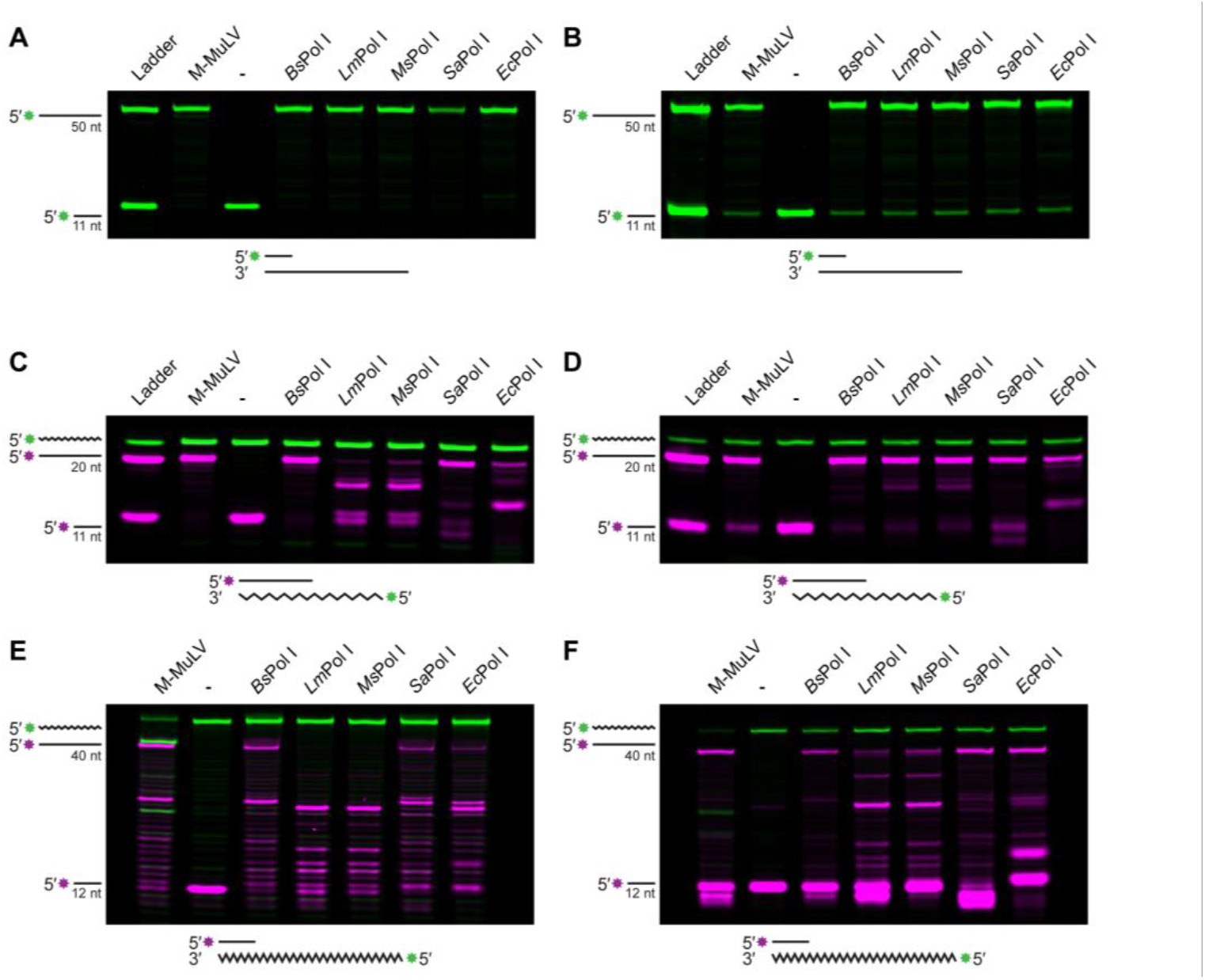
RNA-templated DNA repair is conserved in bacterial Pol I enzymes. (A-C) Primer extension using a 50 nt DNA template (A), a 20 nt RNA template (B) or a 40 nt RNA template using a buffer optimized for a DNA extension. (D-F) The same as A-C, except reactions used a buffer optimized for reverse-transcriptase activity.

Previous work investigating *Ec*Pol I activity using an RNA template has led to inconsistent results, with some groups showing activity^57–61^ and other groups finding activity after protein modification or in the presence of high concentrations of Mn^2+37,57^. Under our assay conditions, *Ec*Pol I was capable of RNA-templated DNA synthesis (Fig. 3). Since one of the papers used commercially available *Ec*Pol I-Klenow fragment and observed no RT activity^37^, we compared the activity of commercially available *Ec*Pol I to the protein assayed in this work and found that both produced similar results under our assay conditions (Sup. Fig. 2). Therefore, we conclude that *Ec*Pol I is capable of RNA-templated DNA synthesis, although the enzyme is not as efficient as *Bs*Pol I or *Sa*Pol I.

### Pol I can traverse embedded ribonucleotides

Since all Pol I enzymes studied here are capable of catalyzing at least some RNA-mediated DNA synthesis, we chose to evaluate their ability to replicate a substrate that contained embedded ribonucleotides, as a model for a relevant circumstance encountered during DNA replication in vivo. We designed a series of substrates with a 15 nt primer-binding site followed by a 5 base “running start” of DNA. After the running start, Pol I would encounter 1, 5, 10 or 15 embedded rNMPs followed by another stretch of DNA (Fig. 4A). Both *Bs*Pol I and *Ec*Pol I could generate full-length products using all templates tested in both the Ext. and RT buffers (Fig. 4B, Sup. Fig. 3A), though the amount of intermediate length product increased for each Pol I as the number of ribonucleotides increased, indicating both Pol Is lose efficiency over a longer patch of rNMPs. We chose to perform this assay for 20 minutes rather than 60 minutes because we found that *Ec*Pol I generates full-length substrate followed by cleavage using its 3′ to 5′ exonuclease activity within the extended timeframe (Sup. Fig. 4). As Figure 4C shows, the other Pol I homologs were not as robust across the different substrates as *Bs* and *Ec*Pol I, although they are clearly capable of traversing short patches of ribonucleotides. As before, the buffer composition impacted efficiency, as the various Pol Is produced more full-length product in the RT buffer than the DNA extension buffer (Fig. 4C, Sup. Fig. 3B).

**Figure 4.**
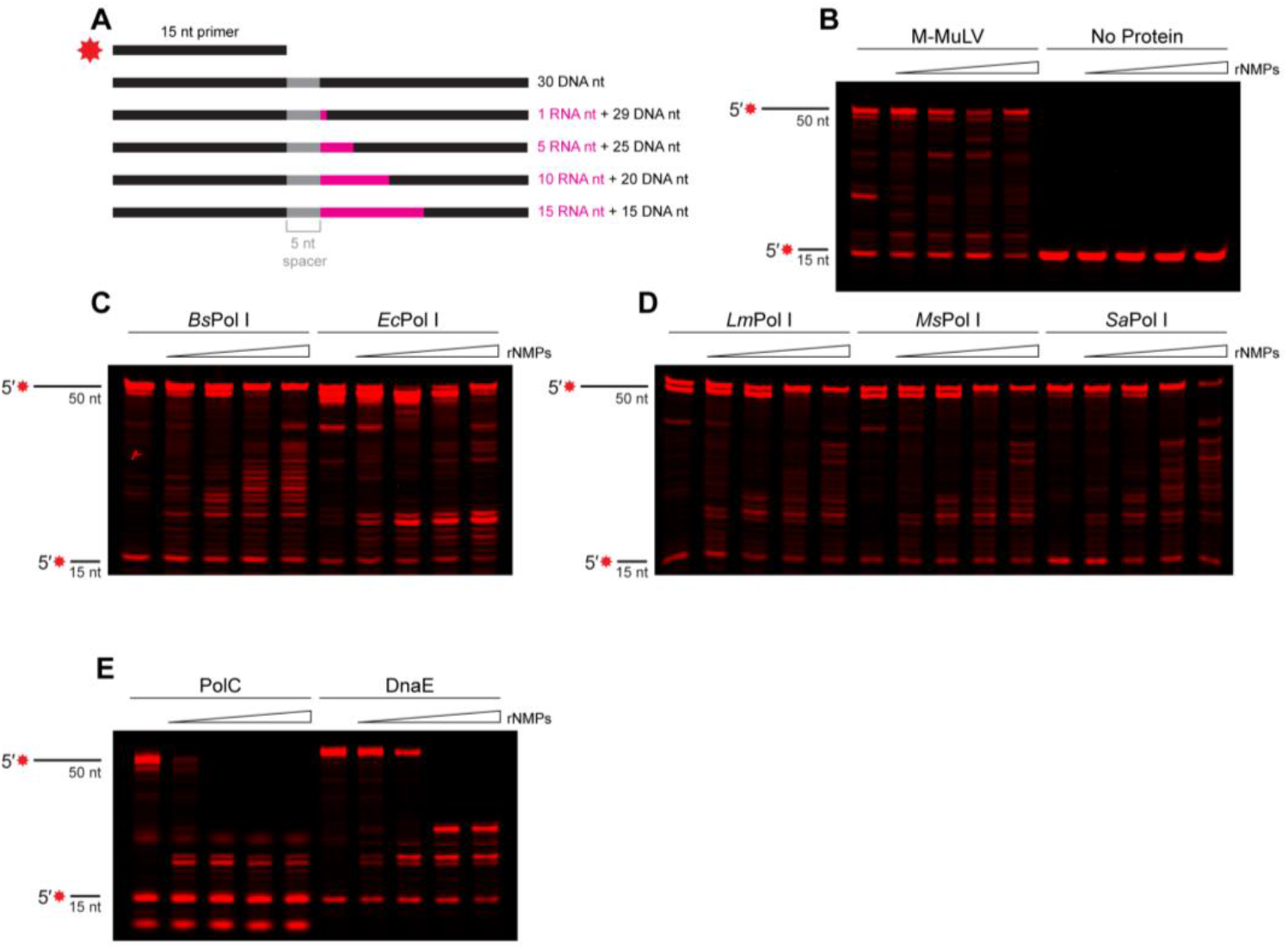
Pol I can traverse embedded ribonucleotides. (A) Schematic of substrates used to mimic ribopatches in DNA. (B-E) Primer extension products generated by the indicated polymerase after 20 minutes using substrates with increasing stretches of ribonucleotides. Assay performed with reverse transcriptase buffer.

While replicative polymerases PolC and DnaE do not have innate reverse transcriptase activity, we asked what number of template ribonucleotides would halt their synthesis activity. PolC was able to fully extend the primer when the template was entirely composed of DNA or contained only one ribonucleotide (Sup. Fig. 3D). The ability to bypass this single rNMP required the specialized Ext. Buffer and was minimal in the more general RT buffer (Fig. 4E). PolC was unable to fully synthesize product when five or more ribonucleotide stretches were present in the template, demonstrating that embedded patches of rNMPs arrest replicative DNA polymerase PolC (Fig. 4D, Sup. Fig. 3E). Interestingly, we did find that DnaE was capable of traversing 1 or 5 rNMPs in the template DNA, however, longer stretches of ribonucleotides did arrest DnaE synthesis (Fig. 4D, Sup. Fig. 3E).. We conclude that Pol I is efficient at RNA-templated DNA synthesis while DnaE is capable of tolerating short patches up to five nucleotides. This suggests that DnaE and Pol I provide two avenues to address 5 or fewer rNMPs in template DNA, with Pol I representing the most efficient option (Fig. 4, Sup. Fig. 3). Further, our data show that the major leading and lagging strand replicative polymerase PolC experiences a nearly complete arrest when more than one rNMP is in the template, demonstrating this polymerase is refractory to RNA-templated DNA repair.

## Discussion

In this work, we demonstrate that DNA polymerase I from *B. subtilis* exhibits robust synthesis activity using RNA-containing templates. This activity allows Pol I to synthesize over embedded ribonucleotides to generate structures recognizable by RNase HII or HIII^50^, which provides cells with another chance to repair nested RNA before threatening genome stability. This would be especially important following instances when intracellular dNTP pools are reduced, as such conditions increase the likelihood of rNMP incorporation by replicative or translesion polymerases^24,62^. Additionally, RNA-templated synthesis would enable Pol I to participate in RNA-mediated double strand break repair^31,37,63^. We validate that the reverse transcriptase activity is relevant to cells, showing that the loss of Pol I increases the hydroxyurea sensitivity of cells lacking RNase HIII, a genotype that is prone to excessive R-loop formation^52^. This demonstrates that Pol I provides a pathway for overcoming RNA-DNA hybrids when they are left unresolved. Previous work suggests that the major role of *Bs*Pol I is DNA repair ^64–67^ rather than Okazaki fragment maturation^68^, consistent with our findings that Pol I participates in RNA-templated DNA synthesis. As such, we suggest that a major role of Pol I is in traversing embedded ribonucleotides encountered by the replication fork.

In contrast, neither PolC nor DnaE can synthesize DNA using an RNA template. When the rNMPs were embedded within a DNA strand, PolC could weakly traverse a single rNMP but consecutive rNMPs arrested synthesis. Given the high incidence of single ribonucleotide misincorporations, and the pausing of PolC at rNMPs^24^, this tolerance demonstrates the necessary balance between processivity and accuracy. Surprisingly, we discovered that DnaE is capable of synthesizing DNA from a template containing a stretch of 5 rNMPs. This suggests that DnaE is sufficient for synthesizing over short stretches of ribonucleotides, but not for longer stretches of incorporated ribonucleotides due to integration from R-loops. This limitation highlights the importance of Pol I’s significant reverse transcriptase activity for addressing broad sources of RNA-DNA hybrids in template DNA. Further, since DnaE lacks 3′ to 5′ proofreading activity, we do find a correlation between a lack of proofreading activity and RNA-templated DNA synthesis, although this is just a correlation and is not absolute as *E. coli* Pol I has proofreading activity and contains RT activity.

We also show that, while the RNA-templated synthesis activity of *Bs*Pol I is generally shared among bacterial Pol Is, *Bs*Pol I performs robustly under remarkably different assay conditions. The other Pol Is we investigate show enhanced activity in a buffer optimized for RT activity, an observation that offers some explanation as to the inconsistent characterization of the RT activity of *Ec*Pol I. Assay conditions vary across prior work, which mean some studies have reported minimal RT activity only under high Mn^2+^ conditions or after protein mutagenesis^37,60^ while others report activity in the absence of these factors^58,59,61^. The consistently high activity of *Bs*Pol I indicates that this polymerase is particularly amenable to non-canonical templates, which is beneficial for a protein whose major role is in DNA repair.

The conservation of reverse transcriptase activity across diverse Pol Is, regardless of their proofreading capabilities, suggests that this function is more broadly modulated. While one group showed different extension from an RNA template using *Ec*PolA-Klenow and *Ec*PolA-Klenow^exo-37^, we find RNA-templated synthesis by the full-length version of *Ec*Pol I. The differences between our work and the prior study likely reflect differences in experimental conditions. Our results do show that the efficiency of the RNA-templated synthesis varies across organisms, and that *Bs*Pol I is an overlooked candidate for determining the structural basis of such disparity. As there are many commercial applications for a DNA polymerase capable of DNA/RNA mediated synthesis, most efforts to elucidate the elements affecting RT activity of these polymerases have focused on improving the activity. In pursuit of such modifications, it has been suggested that the Q-helix, which is a part of the polymerase “fingers” that interact with the template strand^69^ or select residues of the “thumb” domain^70^ may contribute to the adaptability of Pol Is active site. Since A-family polymerases are hypothesized to have evolved from reverse transcriptases^71–73^, differences in active site flexibility may represent broader evolutionary adaptations from a common origin.

In summary, our results demonstrate that Pol I is capable of RNA-templated DNA synthesis and provide evidence that this activity contributes to genome stability under conditions that increase the occurrence of RNA-DNA hybrids. We show that this activity is not shared by the main replicative polymerases, emphasizing the role of A-family polymerases in addressing synthesis over embedded ribonucleotides. These findings highlight the importance of diversity in dealing with a category of damage as broad as RNA-DNA hybrids.

## Acknowledgements

We would like to thank Dr. Charles McHenry for the purified PolC and DnaE used in this work. We would also like to thank Dr. Taylor Nye for providing the genomic DNA used to amplify the *SapolA* gene and Jessica Panchaud for help during the beginning of the project. We thank Lia Munson for contributing to *Bs*Pol I protein purification. This work was funded by National Institutes of Health grant R35GM131772 to LAS.

## Methods

### Spot Titers

*B. subtilis* strains (Supplemental Table 2) were streaked onto LB plates and incubated overnight at 37°C. For each strain, 2 mL of LB was inoculated with a single colony then grown at 37°C in a rolling rack until A_600nm_ was between 0.5 and 0.7. Cultures were pelleted by centrifugation (4000 xg/5 min/25°C) and the supernatant was removed and replaced with 0.85% w/v NaCl (saline). Samples were normalized to A_600nm_ = 0.5, then serially diluted to 10^-5^ in saline. 4 µL of each dilution was plated on LB agar with or without 10 mM hydroxyurea (HU), then grown for 12 hours at 37°C. Spot titers were performed in biological triplicate and plates were prepared on the day of the experiment to minimize HU degradation^49^. Image brightness and contrast were adjusted in FIJI^74^.

### Plasmid Cloning

All primers used are listed in Supplemental Table 3.

The protein sequences for LmPolA (Uniprot ID: A0A6Y7VQD1) and MsPolA (Uniprot ID: I7G3P9) were submitted to Twist Bioscience for gene synthesis and insertion into the pTwist vector harboring Kan resistance. Inserted genes were codon-optimized for expression in E. *coli* and, to simplify downstream cloning, 25 bp of the pE-SUMO vector was added upstream and downstream of the gene.

Ec*polA* (Uniprot ID: P00582) and Sa*polA* (Uniprot ID: A0A0H2XGR4) were amplified using genomic DNA from *E. coli* BL21(DE3) or *S. aureus* USA300 (genomic DNA from Dr. Taylor Nye) as appropriate, while *LmpolA* and *MspolA* were amplified from their respective pTwist plasmids. Empty, linear pE-SUMO expression vector was amplified using oJR46 and 47.

After gel-extraction, each insert was combined with vector using the NEB HiFi protocol (NEB E2621). Assembly reactions were used to transform chemically competent DH5α cells, recovered at 37°C for 1 hour, and grown overnight at 37°C on LB agar supplemented with 25 μg/mL kanamycin. Transformants were screened using colony PCR with prFCL124 and 125, then plasmid sequence was confirmed with full-plasmid sequencing.

### Protein Expression

Chemically competent *E. coli* BL21(DE3) were transformed with plasmid (Supplemental Table 4) and plated on LB+Kan (25 μg/mL kanamycin, unless stated otherwise) and grown overnight at 37°C. Starter culture was prepared by inoculating LB+Kan with transformant cells, which was then grown overnight at 37°C while shaking. For expression, each liter of LB+Kan was inoculated with starter culture, then grown to A_600nm_ ~ 0.7. Cultures were induced by adding iso-propyl ß-D-1-thiogalactopyranoside (IPTG) to 0.5 mM, then grown for 3 hours at 37°C. Induced cells were harvested by centrifugation at 4000 xg /4°C/25 min, after which the supernatant was discarded, and pellets were stored at −80°C.

### Protein Purification

All purification steps were performed at 4°C. Protein pellets were thawed on ice, then resuspended in lysis buffer (20 mM Tris pH 8.0, 400mM NaCl, 5 % v/v glycerol), supplemented with protease inhibitor (Pierce: A32965) and 1 mM phenylmethanesulfonyl fluoride (PMSF). Cells were lysed via sonication, then the lysate was clarified by centrifugation at 30000 xg for 30 min. Clarified lysate was applied to a gravity column containing Ni^2+^-NTA-agarose that had been pre-equilibrated in lysis buffer and allowed to pass through. The column was washed with 5 CV lysis buffer, then with 5 CV of wash buffer (20 mM Tris pH 8.0, 2 M NaCl, 15mM imidazole, 5 % v/v glycerol). Protein that remained bound to the column was eluted with 3 CV elution buffer (20 mM Tris pH 8.0, 400 mM NaCl, 300 mM imidazole, 5 % v/v glycerol). The wash and elution fractions were evaluated using SDS-PAGE, then the protein-containing fractions were pooled and treated with Ulp1-protease for 2 hours at room temperature to remove the 6x-His-SUMO tag. The protein was then dialyzed overnight against Dialysis Buffer (20 mM Tris pH 8.0, 300mM NaCl, 5 % v/v glycerol). Dialyzed protein was then passed over Ni^2+^-NTA-agarose that had been equilibrated with dialysis buffer. The column was washed with 3 CV dialysis buffer and all flowthrough was collected and diluted to 50 mM NaCl with Q buffer A (20 mM Tris pH 8.0, 5 % v/v glycerol, 1 mM DTT). This protein was filtered with a 0.22 µm syringe filter and applied to a HiTrap Q FF anion exchange column (Cytivia, 17515601) and fractionated with an AKTA FPLC. Protein was eluted using an increasing gradient of Q buffer B (20 mM Tris pH 8.0, 500 mM NaCl, 5 % v/v glycerol, 1 mM DTT). Peak fractions were evaluated using SDS-PAGE, then pooled and concentrated using a centrifugal filter. Glycerol was added to 25%, then proteins were aliquoted, flash-frozen, then stored at −80°C. The gel demonstrating the proteins used in this paper was made by loading 2 µg of each protein onto a Mini-PROTEAN TGX 4–20% gradient gel (BIO-RAD, 4561096) and then staining with Coomassie blue.

### Extension Assays

Extension assays were carried out as described previously^53,68^. In brief, annealed oligonucleotide substrates were generated by combining 1 µM of the primer and 2 µM of the template in 1x Dilution Buffer (20 mM Tris, pH 8.0, 25 mM NaCl) then heating them at 98°C for 1 minute. Substrates were allowed to cool to room temperature. All oligonucleotides used were purchased from IDT and sequences are included in Supplemental Table 2. Substrates were made using the following oligonucleotide combinations. The DNA templates used in Figures 2 and 3 use oFCL7 and 8, with oFCL27 as the ladder. The short RNA templates used in Figures 2, 3 and Sup. Figure 2 use oFCL16 and oJR227, with oFCL20 as the ladder. The long RNA templates used in Figures 2 and 3 use oFCL18 and oJR336. Embedded ribonucleotide substrates (Figure 4, Sup. Figures 3, and Sup. Fig 4) consist of oFCL21 annealed to oFCL22, 23, 24, 25, or 26.

For each reaction, a new aliquot of protein was diluted to 400 nM in 1x Metals Buffer (20 mM Tris, pH 8.0, 50 nM NaCl, 1 mM MgCl_2_). Protein was added, at a final concentration of 100 nM, to a reaction mix containing 100 nM substrate, 50 µM deoxynucleotide triphosphates (dNTPs) and 1x Extension Buffer (40 mM Tris-acetate,pH 7.8, 12 mM magnesium acetate, 300 mM potassium glutamate, 3 μM ZnSO4, 2% (w/v)polyethylene glycol, 0.02% pluronic F68) or 1x Reverse Transcription Buffer (RT Buffer) (50 mM Tris pH 8.0, 75 mM KCl, 3 mM MgCl_2_). The RT Buffer was based on the 1X M-MuLV Reverse Transcriptase Reaction Buffer (New England Biolabs, B0253S). M-MuLV (M0253S) and *E. coli* Pol I (M0209S) controls were purchased from New England Biolabs. Commercial protein concentrations were estimated using Nanodrop spectrophotometry and were used at the same concentrations as the other proteins, which translates to 2.4 units of M-MuLV and 0.15 units of *Ec*Pol I per reaction. Assays were performed at 37°C for 60 minutes, except for those using the embedded ribonucleotide substrates, which were incubated at 37°C for 20 minutes (Fig. 4) or for the time indicated (Sup. Fig. 4). Reactions were terminated by adding an equivalent volume of Stop Buffer (95% formamide, 20nM EDTA, bromophenol blue) to the reaction, boiling them for 10 minutes at 98°C, then snap-cooling in ice. Reaction products were resolved by Urea-PAGE using gels containing 20% Acrylamide and 8M urea. To minimize the persistence of nucleic acid complexes, the 1x TBE running buffer was pre-warmed to approximately 45°C and the electrophoresis box was kept on a 45°C hot plate while voltage was applied. Gels were visualized using the 700 and 800 nm channels of a LiCor Oddesey CLx, and channel color was altered from red/green to magenta/green using FIJI^74^. Original colors are included in Sup. Table 2.

## Notes

### Competing Interest Statement

The authors have declared no competing interest.

