## Supplemental Tables S1-4 and Figures S1-4 for "DNA polymerase I is an efficient reverse transcriptase that mediates RNA-templated DNA repair synthesis"

Short title:

**Keywords:** Reverse transcriptase, *Bacillus subtil*is, RNA-DNA hybrid, genome instability, DNA polymerase I

**Supplemental Table 1.** Bacterial Pol Is with reverse transcriptase activity

| Organism | Gram-stain | D | E | D | Y-H | D | Active 3′-5′ Exonuclease? | Notes |
|---|---|---|---|---|---|---|---|---|
| *Bacillus caldolyticus* EA.1[1] | Positive | - | E | - | H | - | No | |
| *Bacillus subtilis* | Positive | - | E | - | H | - | No | First studied in this work. |
| *Caldibacillus cellulovorans* CompA.2[1] | Positive | - | E | - | Y | - | No | |
| *Caldicellulosiruptor saccharolyticus* Rt69B.1[1] | Positive | - | - | - | - | - | No | |
| *Caldicellulosiruptor saccharolyticus* Tok13B.1[1] | Positive | - | - | - | - | - | No | |
| *Caldicellulosiruptor saccharolyticus* Tok7B.1[1] | Positive | - | - | - | - | - | No | |
| *Dictyoglomus thermophilum* Rt46B.1[1] | Negative | - | - | D | H | - | No | |
| *Escherichia coli*[2–8] | Negative | D | E | D | Y | D | Yes | |
| *Geobacillus stearothermophilus*[9–11] | Positive | - | E | - | H | - | Yes | |
| *Listeria monocytogenes* | Positive | - | E | - | H | - | Yes | First studied in this work. |
| *Micrococcus luteus*[3] | Positive | - | - | - | - | - | No | |
| *Mycolicibacterium smegmatis*[12] | - | - | - | - | - | - | No | Bound 1 $Mn^{2+}$ using non-conserved residues[12] |
| *Pseudomonas aeruginosa*[7] | Negative | D | E | D | Y | D | Yes | |
| *Shigella sonnei*[7] | Negative | D | E | D | Y | D | Yes | |
| *Staphylococcus aureus*[7] | Positive | - | E | - | Y | - | No | |
| *Streptococcus agalactiae*[13] | Positive | - | E | - | H | - | No | |
| *Streptomyces coelicolor*[14] | Positive | - | - | - | - | - | No | |
| *Thermoactinomyces vulgaris*[9] | Positive | - | E | - | H | - | No | |
| *Clostridium thermosulfurogenes*[1] | Positive | - | - | - | Y | - | No | |
| *Thermoclostridium stercorarium*[1] | Positive | - | - | - | Y | - | No | |
| *Thermotoga neapolitana*[1,15] | Negative | D | E | D | Y | D | Yes | Modified to lose 3′-5′[1] |
| *Thermus aquaticus*[1,16,17] | Negative | - | - | - | - | - | No | |
| *Thermus filiformis*[1] | Negative | - | - | - | - | - | No | |
| *Thermus thermophilus*[1,18,19] | Negative | - | - | - | - | - | No | Activity studied with $Mn^{2+}$[18,19] |
| *Vibrio cholerae* non-O1[7] | Negative | D | E | D | Y | D | Yes | |

Note that the identity of the *E. coli* hybrid polymerase purified by Lee-Huang and Cavalieri is unclear[6], but the reference is included in Sup. Table 1 since other works studying *Ec*Pol I make reference to it[4].

**Supplemental Table 2.** *B. subtilis* strains used in this study.

| Strain Identifier | Relevant Genotype | Citation |
|---|---|---|
| FCL3 | ∆*polA* | JWS235[20] |
| FCL10 | native PY79 | Youngman |
| FCL11 | ∆*rnhC* | JRR48[20] |
| FCL12 | ∆*rnhC*, ∆*polA* | JRR64[20] |

**Supplemental Table 3.** Oligonucleotides used in this study.

| Oligonucleotide | Purpose | Sequence (5′-3′) |
|---|---|---|
| oJR46 | Amplifying pE-SUMO vector (F) | TCGAGCACCACCACCACCACCACTGAG |
| oJR47 | Amplifying pE-SUMO vector (R) | ACCTCCAATCTGTTCGCGGTGAGCCTCAATAATATCG |
| prFCL120 | Amplifying *SapolA* (F) | ctcaccgcgaacagattggaggtGTGAATAAATTAGTATTAATCGATGG |
| prFCL121 | Amplifying *SapolA* (R) | gtggtggtggtggtgctcgaTTATTTTGCATCATACCAGGTTGCACC |
| prFCL122 | Amplifying Ec*polA* (F) | ctcaccgcgaacagattggaggtATGGTTCAGATCCCCCAAAATCCACTTATCC |
| prFCL123 | Amplifying Ec*polA* (R) | gtggtggtggtggtgctcgaTTAGTGCGCCTGATCCCAGTTTTCG |
| prFCL124 | Amplifying pTwist Insert (F) | GGCTCACCGCGAACAGATTGGAGGT |
| prFCL125 | Amplifying pTwist Insert (R) | CAGTGGTGGTGGTGGTGGTGCTCGA |
| oFCL7 | Assay – DNA Template | GCA*A*T*CGACTCGTAAGCATGGTTCACTACTCGCTGCTTGATGCTCAATCG |
| oFCL8 | Assay – DNA Primer | /5IRD800/C*G*A*TTGAGCATCAAGCAGCG |
| oFCL16 | Assay – Short RNA Primer | /5IRD700/G*C*A*GAGCTAGC |
| oFCL18 | Assay – Long RNA Primer | /5IRD700/A*C*A*GCGTTCCCT |
| oFCL20 | Assay – Short RNA Ladder | /5IRD700/GCAGAGCTAGCTTACGATCG |
| oFCL21 | Assay – Hybrid Primer | /5IRD700/C*A*A*GTCATCAAATGG |
| oFCL22 | Assay – Hybrid DNA Template | TGAGTAAGTTGGTATCCGAGGTACTATGAGCTTCTGGACCATTTGATGACTTG |
| oFCL23 | Assay – Hybrid 1 nt Template | TGAGTAAGTTGGTATCCGAGGTACTATGAGCTUCTGGACCATTTGATGACTTG |
| oFCL24 | Assay – Hybrid 5 nt Template | TGAGTAAGTTGGTATCCGAGGTACTATGAGCUUCTGGACCATTTGATGACTTG |
| oFCL25 | Assay – Hybrid 10 nt Template | TGAGTAAGTTGGTATCCGAGGTACUAUGAGCUUCTGGACCATTTGATGACTTG |
| oFCL26 | Assay – Hybrid 15 nt Template | TGAGTAAGTTGGTATCCGAGGUACUAUGAGCUUCTGGACCATTTGATGACTTG |
| oFCL27 | Assay – DNA Ladder | /5IRD800/C*G*A*TTGAGCATCAAGCAGCGAGTAGTGAACCATGCTTACGAGTCGATTGC |
| oJR227 | Assay – Short RNA Template | /5IRD800CWN/CGAUCGUAAGCUAGCUCUGC |
| oJR336 | Assay – Long RNA Template | /5IRD800CWN/CUGGAGGAUGGAGGAUGGUGGAGAUGUGAGGGAACGCUGU |

* indicates phosphorothioate linkages

IRD indicates an infrared dye at the 5′ end or 3′ end as indicated, 700 or 800 indicates the wavelength of excitation, and CWN indicates a NHS ester conjugation.

Red bases indicate ribonucleotides.

**Supplemental Table 4.** Plasmids used in this study.

| **Plasmid Identifier** | **Vector** | **Insert** |
|---|---|---|
| pJR22 | pE-SUMO | *BspoIA*[21] |
| pFCL56 | pE-SUMO | *SapolA* |
| pFCL57 | pTwist-Kan | *LmpolA** |
| pFCL58 | pTwist-Kan | MspolA* |
| pFCL59 | pE-SUMO | *EcpolA* |
| pFCL61 | pE-SUMO | *LmpolA** |
| pFCL62 | pE-SUMO | MspolA* |

***** indicates genes codon optimized for expression in *E. coli*

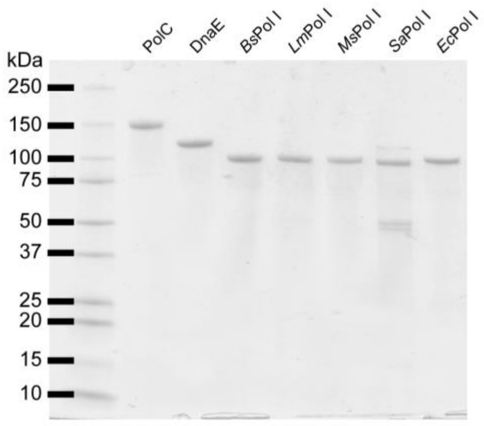


**Sup. Figure 1. Purified proteins used in biochemical assays.** 2 µg of each protein used for extension assays are shown on a Coomassie-stained, SDS-PAGE.

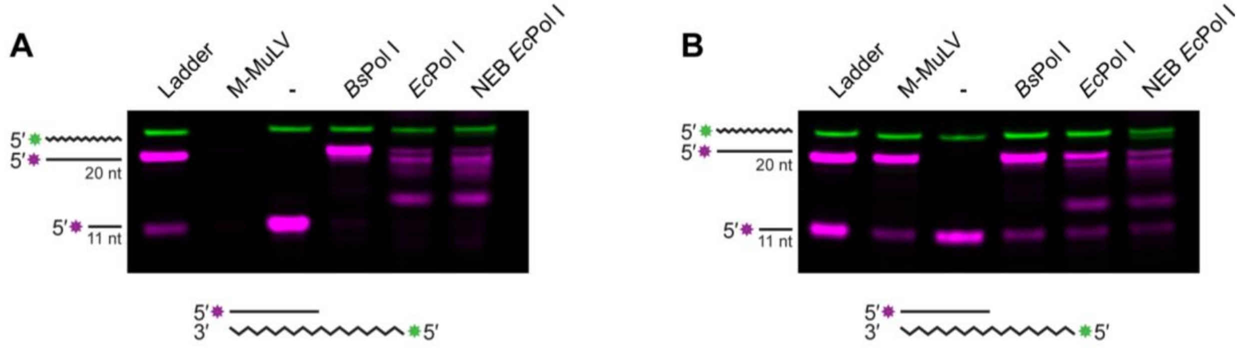


**Sup. Figure 2. Commercial EcPol I has reverse transcriptase activity.** (A) Primer extension in DNA extension buffer using an RNA template. (B) Primer extension from an RNA template in reverse transcriptase buffer.

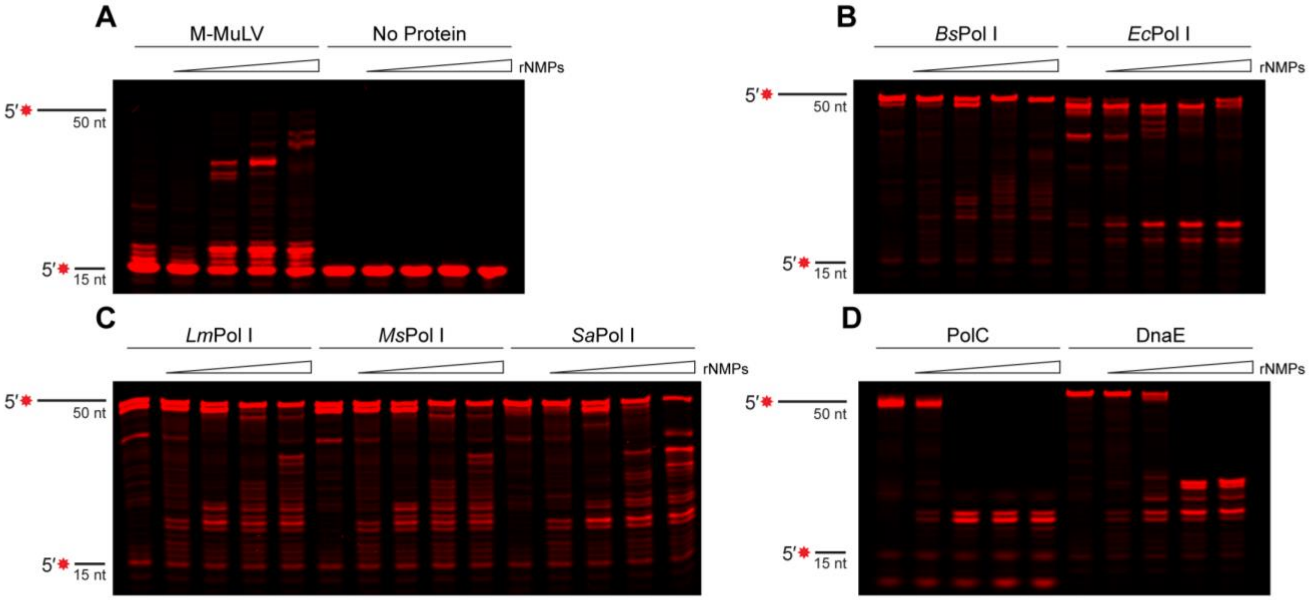


**Sup. Figure 3. Pol I traverses embedded ribonucleotides under DNA extension conditions.** (A-D) Primer extension products generated by the indicated polymerase after 20 minutes using substrates with increasing stretches of ribonucleotides.

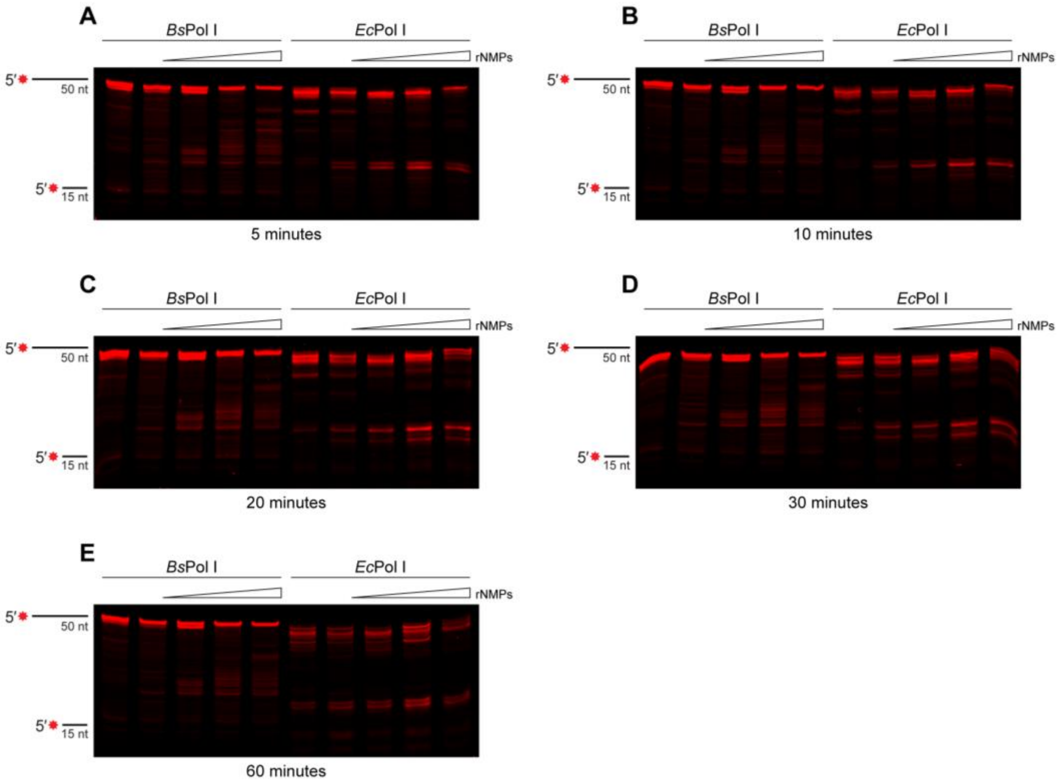


**Sup. Figure 4. EcPol I nuclease activity degrades reaction products over time.** (A-E) Primer extension products generated by the indicated polymerase using substrates with increasing stretches of ribonucleotides. Reaction incubation times are indicated below each gel.
